# Updated MS^2^PIP web server delivers fast and accurate MS^2^ peak intensity prediction for multiple fragmentation methods, instruments and labeling techniques

**DOI:** 10.1101/544965

**Authors:** Ralf Gabriels, Lennart Martens, Sven Degroeve

**Author notes:** To whom correspondence should be addressed, Address: VIB-UGent Center for Medical Biotechnology, A. Baertsoenkaai 3, B9000 Ghent, Belgium.

## Abstract

MS^2^PIP is a data-driven tool that accurately predicts peak intensities for a given peptide’s fragmentation mass spectrum. Since the release of the MS^2^PIP web server in 2015, we have brought significant updates to both the tool and the web server. In addition to the original models for CID and HCD fragmentation, we have added specialized models for the TripleTOF 5600+ mass spectrometer, for TMT-labeled peptides, for iTRAQ-labeled peptides, and for iTRAQ-labeled phosphopeptides. Because the fragmentation pattern is heavily altered in each of these cases, these additional models greatly improve the prediction accuracy for their corresponding data types. We have also substantially reduced the computational resources required to run MS^2^PIP, and have completely rebuilt the web server, which now allows predictions of up to 100,000 peptide sequences in a single request. The MS^2^PIP web server is freely available at https://iomics.ugent.be/ms2pip/.

## INTRODUCTION

In high throughput tandem mass spectrometry (MS^2^), peptides are identified by analyzing their fragmentation spectra. These spectra are obtained by collision induced dissociation (CID) or higher-energy collisional dissociation (HCD), where peptides are made to collide with an inert gas, or by electron-transfer dissociation (ETD) or electron-capture dissociation (ECD), in which electrons are transferred to peptides. After fragmentation, the mass-to-charge ratios (m/z) and intensities of the resulting fragment ions are measured, yielding the two dimensions of a fragmentation spectrum. While the fragment ions’ m/z can easily be calculated for any given peptide, their intensities have proven to follow extremely complex patterns (1).

In 2013, we therefore developed the data-driven tool MS^2^PIP: MS^2^ Peak Intensity Prediction (2), which can predict fragment ion intensities. By applying machine learning algorithms on the vast amounts of data present in public proteomics repositories such as the PRIDE Archive (3, 4), we could create generalized models that accurately predict the expected normalized MS^2^ peak intensities for a given peptide. While the first iteration of MS^2^PIP outperformed the then state-of-the art prediction tool PeptideART (5), it was originally only trained for CID fragmentation spectra. As HCD fragmentation became more popular in the field, we therefore expanded MS^2^PIP with prediction models for HCD spectra. In 2015, we built the MS^2^PIP web server to make these models easily available to all potential users, regardless of their computational resources (6).

Over the past few years, MS^2^PIP has been used by researchers to create proteome-wide spectral libraries for proteomics search engines (including Data Independent Acquisition), to select discriminative transitions for targeted proteomics (7, 8), and to validate interesting peptide identifications (e.g. biomarkers) (9, 10). Moreover, we have also shown that MS^2^PIP predictions can be used to improve upon and even replace proteomics search engine output when rescoring peptide-to-spectrum matches (11).

Because of the great interest in, and steadily increasing relevance of, MS^2^ peak intensity prediction, we have continued to update and improve MS^2^PIP and the MS^2^PIP web server. We have updated MS^2^PIP to be more computationally efficient, we have rebuilt the MS^2^PIP web server to handle up to 100,000 peptide sequences per request instead of 1,000, and we have added specialized models for the TripleTOF 5600+ mass spectrometer and for isobaric labeled peptides.

## NEW IN THE 2019 VERSION OF MS^2^PIP

### More efficient MS^2^PIP code

Rapid advances in machine learning research combined with larger and more diverse training datasets have allowed for more accurate MS^2^PIP predictive models. The Random Forest algorithm employed in the original MS^2^PIP has made room for a Gradient Tree Boosting algorithm (12), which, in combination with more training data, has improved prediction accuracy. This improved prediction is especially noticeable for peptides with higher charge states, where the large performance differences between charge 2+ and 3+ observed for the original MS^2^PIP models have been significantly reduced in the new version (Supplementary Figure 1).

In addition, we have drastically reduced the required computational resources for MS^2^PIP, while simultaneously further improving its prediction speed. The large memory footprint of the original version (requiring several gigabytes) has now been reduced to just a few hundred megabytes, depending on input request size. When run locally on a normal four core laptop, MS^2^PIP can predict peak intensities for a million peptides in less than five minutes.

### Specialized models for isobaric labeled peptides and the TripleTOF 5600+ mass spectrometer

One of the most important changes in this new version of MS^2^PIP is the addition of specialized models for specific types of peptide spectra. The type of mass spectrometer, fragmentation method and certain peptide modifications (such as isobaric labels and phosphorylation) can heavily alter peptide fragmentation patterns. We have therefore now also trained specialized models for the TripleTOF 5600+ mass spectrometer, for TMT-labeled peptides (13), for iTRAQ-labeled peptides (14), and for iTRAQ-labeled phosphopeptides (Table 1). Each of these models was trained and evaluated on publicly available spectral libraries or experimental datasets, ranging in size from 183,000 to 1.6 million peptide spectra. Final validation of every model was based on wholly independent datasets, ranging in size from 9,000 to 92,000 unique peptide spectra (Table 2). Spectral libraries were filtered for unique peptides and then converted to MS^2^PIP input format. For experimental datasets, original peptide identifications as provided by the data submitter were used where available. Where such original identifications were not available, we performed the identification using the MS-GF+ (15) search engine in combination with Percolator (16) for post-processing.

**Table 1.** All specialized MS^2^PIP models with MS^2^ acquisition information and peptide properties of the training datasets.

| Model | Fragmentation method | MS<sup>2</sup> mass analyzer | Peptide properties |
| --- | --- | --- | --- |
| CID | CID | Linear ion trap | Tryptic digest |
| HCD | HCD | Orbitrap | Tryptic digest |
| TripleTOF 5600+ | CID | Quadrupole Time-of-Flight | Tryptic digest |
| TMT | HCD | Orbitrap | Tryptic digest, TMT-labeled |
| iTRAQ | HCD | Orbitrap | Tryptic digest, iTRAQ-labeled |
| iTRAQ phospho | HCD | Orbitrap | Tryptic digest, iTRAQ-labeled enriched for phosphorylation |

**Table 2.**
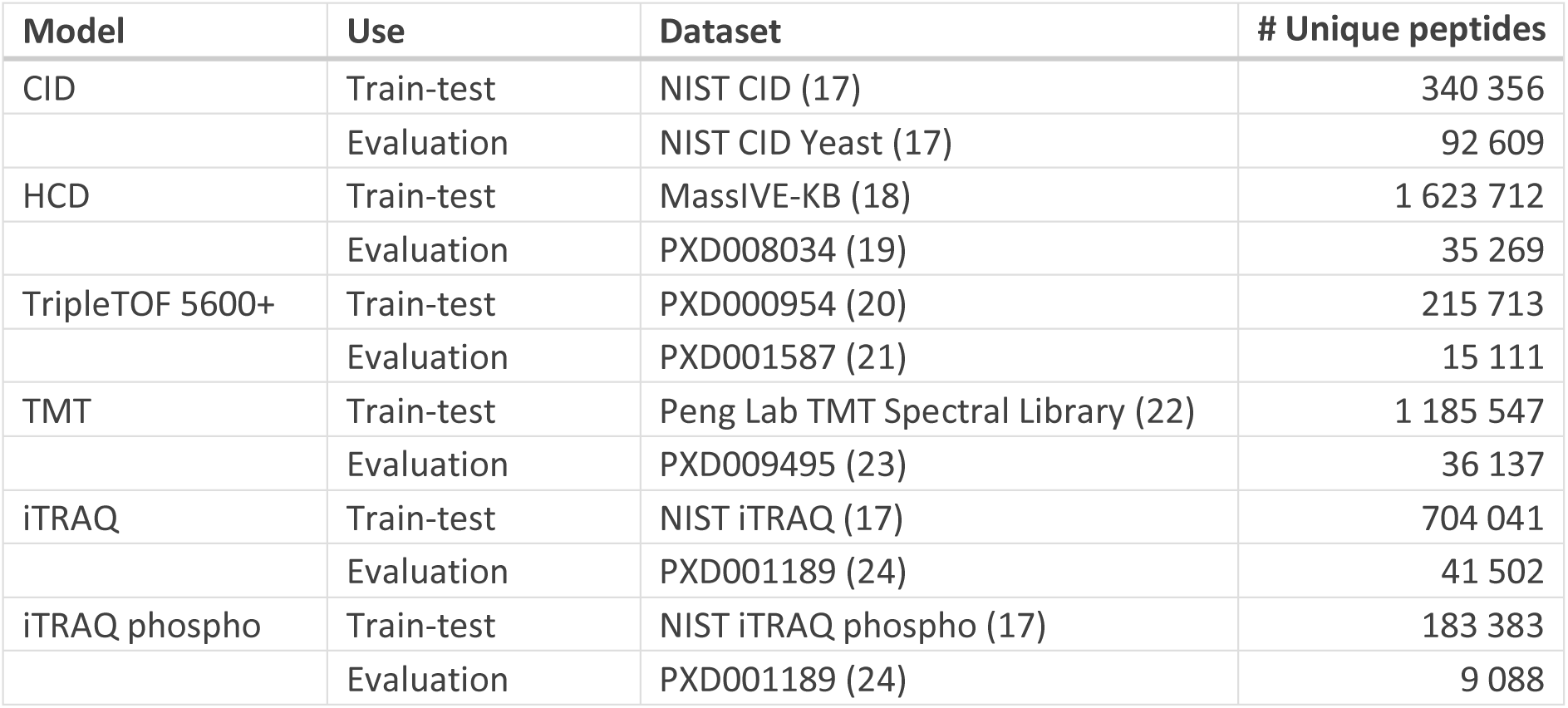
Train-test and evaluation datasets used for specialized MS^2^PIP models.

### Redesigned, more robust web server

Along with the heavily updated MS^2^PIP models, we have also rebuilt the web server from the ground up. Like the previous version, this web server has been built using the Flask framework (https://flask.pocoo.org) with a front-end based on Bootstrap (https://getbootstrap.com).

In this newly built web server, we have implemented a robust queueing system that is able to handle concurrent tasks. This has allowed us to increase the maximum number of peptide sequences *per* request from 1,000 to 100,000. Besides submitting a single task through the website, users can also automate their requests through MS^2^PIP’s updated RESTful API, for which we provide an example Python script. A single request of 100,000 peptide sequences takes less than five minutes to complete, including up- and download time. Predictions for 1,000 peptide sequences are returned in less than three seconds.

On the user-friendly webpage, users can select one of the available models and upload a csv file with peptide sequences, precursor charges, and modifications. After uploading this input file, a progress bar displays the status of the request and a URL is displayed to which the user can return at any time to check the status of their request (e.g., in case the browser window was closed). When the predictions have been finalized, the user can inspect the results through several interactive plots, and the predicted spectra can be downloaded in comma-separated values (CSV) format, in Mascot Generic File (MGF) format, in BiblioSpec or Skyline (SSL and MS2) formats (25, 26), or in NIST (National Institute of Standards and Technology) MSP spectral library format.

## PERFORMANCE OF THE SPECIALIZED MODELS

We can evaluate MS^2^PIP model performance by predicting peak intensities for peptides present in the external evaluation datasets, and by comparing these predictions to their corresponding empirical spectra. This comparison is performed through the Pearson correlation coefficient (PCC) between predicted and experimental spectra. The resulting PCC distributions for each of the specialized models are shown in Figure 1A.

The median PCCs are higher than 0.90 for all models, except for the TripleTOF 5600+ and the iTRAQ phospho models, which have median PCCs of 0.74 and 0.84, respectively. These two lower median correlations might be the result of lower training dataset sizes (see also Table 2).

When we apply all specialized models to each specific evaluation dataset – that is, including mismatched model-dataset combinations, such as applying the TMT model to the HCD evaluation dataset – we consistently observe median PCCs that are substantially higher for correctly matched models and evaluation datasets than for mismatched models and evaluation datasets (Figure 1B). Only the specialized TripleTOF 5600+ model is comparable in performance to the HCD model when predicting TripleTOF 5600+ spectra. Overall, this figure makes a clear case for the utility of specialized MS^2^PIP models for specific types of data.

Figure 1B also shows which specialized cases have similar fragmentation patterns. The specialized models for isobaric-labeled peptides (TMT, iTRAQ, and iTRAQ phospho) are quite similar in performance across the different evaluation datasets, as are the HCD and TripleTOF 5600+ models. To further verify this, we have directly compared the models by calculating the PCCs for all specialized model predictions for the same set of peptides (Supplementary Figure 2). The results confirm the findings we observe in Figure 1.

**Figure 1.**
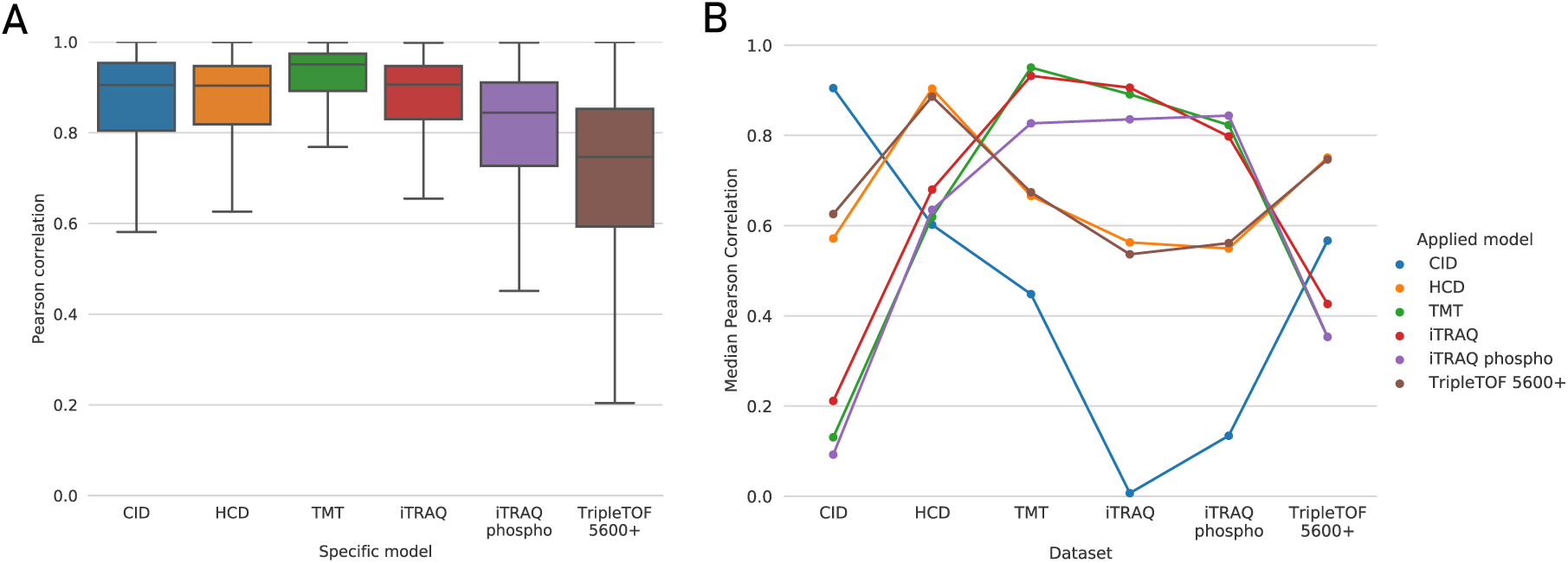
A) Boxplots showing the Pearson correlation coefficients (PCCs) for each of the specialized models applied to their respective evaluation dataset. B) Median PCCs when applying all specialized models to each evaluation dataset, showing the utility of specialized models. Each dot shows the median PCC of a specialized model applied to a specific evaluation dataset. To improve readability, dots representing performance of a single model are connected.

We can also visualize the differences in fragmentation pattern by plotting the predictions from two different models for the same peptide sequence and mirroring the empirical spectrum below these predictions. This is shown in Figure 2 for the TMT and HCD models with an empirical TMT-labeled peptide spectrum. While the TMT model mirrors the empirical TMT spectrum very well, the HCD model does not match the empirical TMT spectrum.

**Figure 2.**
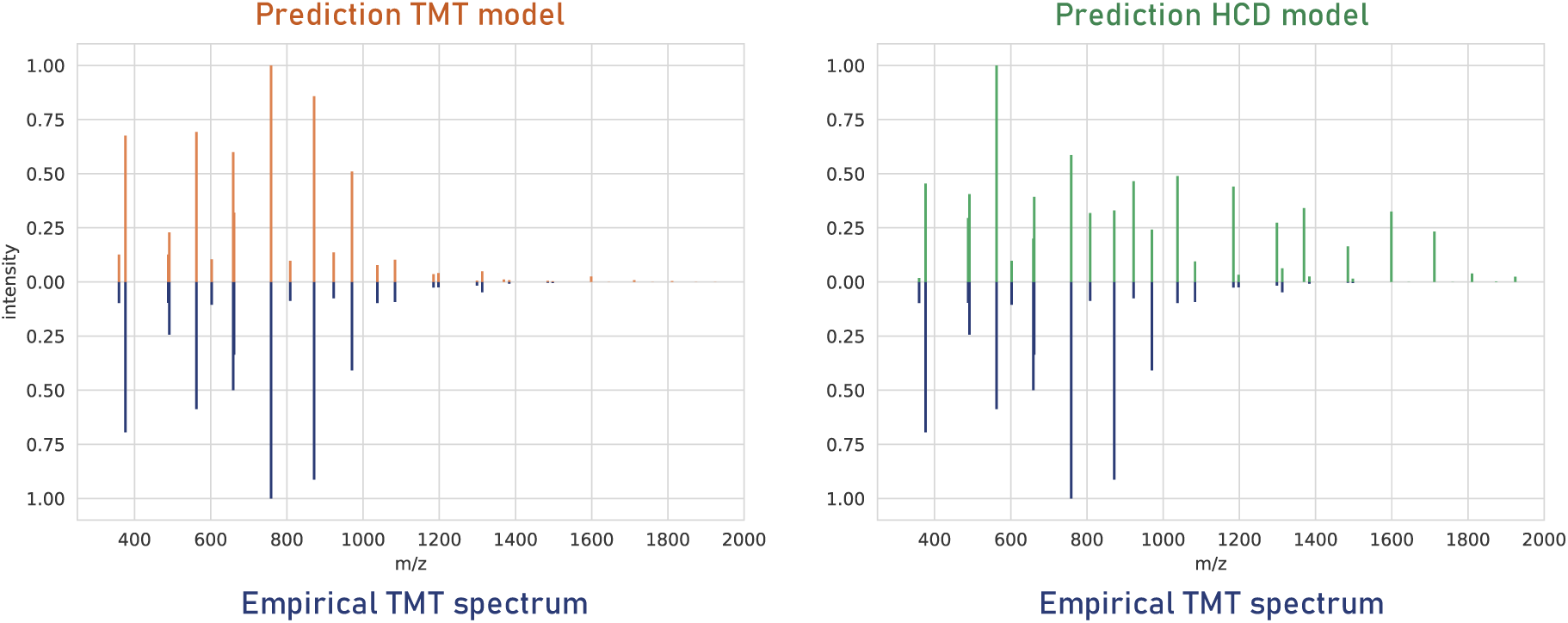
Predictions for the peptide sequence EENGVLVLNDANFDNFVADK, carrying two TMT labels, produced by the TMT model (top left) and the HCD model (top right), compared to the empirical spectrum (bottom left and right).

An additional parameter that influences fragmentation patterns is the collision energy (CE). Yet, as most spectral libraries do not include information on the CE values, CE is not part of MS^2^PIP’s feature set. In order to evaluate MS^2^PIP’s performance across different CEs, we have therefore applied the HCD model on a large public dataset of synthetic peptides measured at different CEs (27). The results are shown in Supplementary Figure 3. For confident PSMs (Andromeda score higher than 200) at higher CE values (30% and 35% normalized CE), median PCCs are above 0.90, which corresponds to the general HCD model evaluation. For confident PSMs at a lower CE value of 25% normalized CE, the median PCC is slightly lower at 0.85. It therefore seems that most real-life data is recorded at higher CE values, as the overall HCD performance of MS^2^PIP most closely resembles 30% and 35% normalized HCD. As the overall HCD performance already indicated, MS^2^PIP will thus produce reliable peak intensity predictions in typical applications. Nevertheless, it is important to be mindful of the effect of altered CE values when interpreting MS^2^PIP predictions, especially in those cases where lower CEs were used.

## CONCLUSION AND FUTURE PERSPECTIVES

With the advent of novel mass spectrometry methods and new computational pipelines, MS^2^ peak intensity prediction is becoming ever more relevant. As one of the front runners in peak intensity prediction, MS^2^PIP has already been used for a variety of purposes, including creation of proteome-wide spectral libraries, optimization of targeted proteomics applications, validation of interesting peptide identifications, and rescoring of search engine output.

With the current update, we present our latest efforts in further widening the scope of MS^2^PIP. The new web server enables researchers to easily obtain more predictions more efficiently, and the new MS^2^PIP models extend the applicability of MS^2^PIP to more varied, popular use cases, allowing it to be applied when specific fragmentation methods, instruments, or labeling techniques are employed.

## AVAILABILITY

The MS^2^PIP web server is freely available via https://iomics.ugent.be/ms2pip. Documentation for contacting the RESTful API is available via https://iomics.ugent.be/ms2pip/api/. MS^2^PIP is open source, licensed under the Apache-2.0 License, and is hosted on https://github.com/compomics/ms2pip_c. All Python scripts that were used to generate the figures are available in a Jupyter notebook via https://github.com/compomics/ms2pip_c/tree/releases/manuscripts/2019.

## ACKNOWLEDGEMENT

We would like to thank all researchers who made their mass spectrometry data publicly available.

