## Supplementary Figures for "Updated MS^2^PIP web server delivers fast and accurate MS^2^ peak intensity prediction for multiple fragmentation methods, instruments and labeling techniques"

Ralf Gabriels[1,2], Lennart Martens[1,2*], Sven Degroeve[1,2]
1. VIB-UGent Center for Medical Biotechnology, VIB, Ghent, Belgium
2. Department of Biomolecular Medicine, Faculty of Medicine and Health Sciences, Ghent University, Ghent, Belgium

* To whom correspondence should be addressed:


Address:
VIB-UGent Center for Medical Biotechnology
A. Baertsoenkaai 3
B9000 Ghent, Belgium

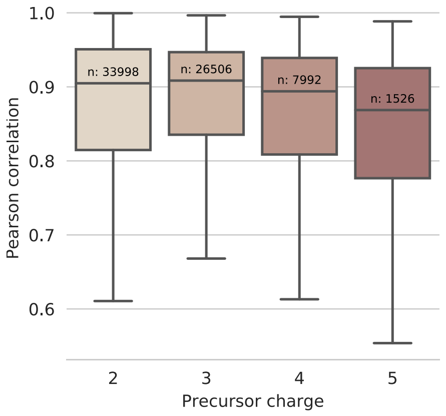


B

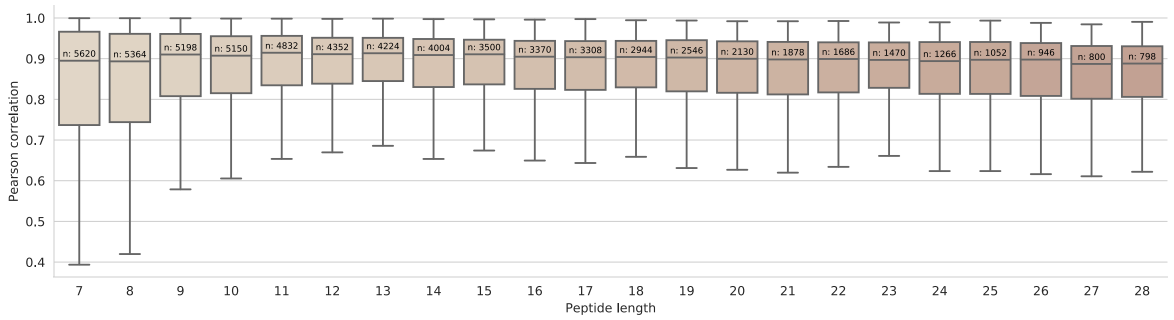


**Supplementary Figure 1.** Boxplots showing the Pearson correlation coefficients for the HCD model applied to the HCD evaluation dataset split by precursor charge (A) and peptide length (B). Only boxplots containing more than 750 datapoints are plotted. The number in each boxplot displays its number of datapoints.

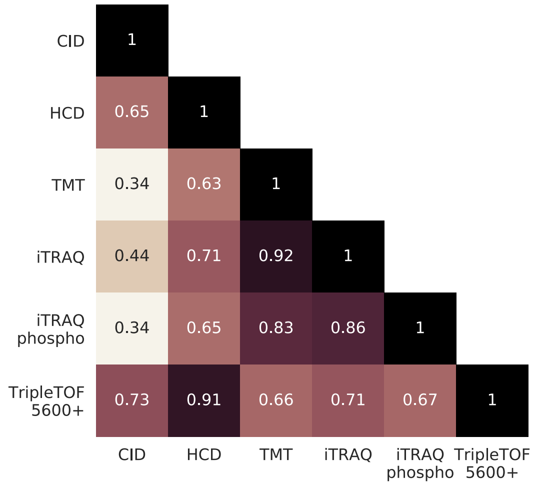


**Supplementary Figure 2.** Correlation matrix directly comparing the different model predictions. Pearson correlation coefficients were calculated between the predictions of all specialized models on a large list of peptides. The numbers in each box correspond to the median Pearson correlation coefficient between the model on the x-axis and the model on the y-axis. A darker color indicates a higher median Pearson correlation coefficient.

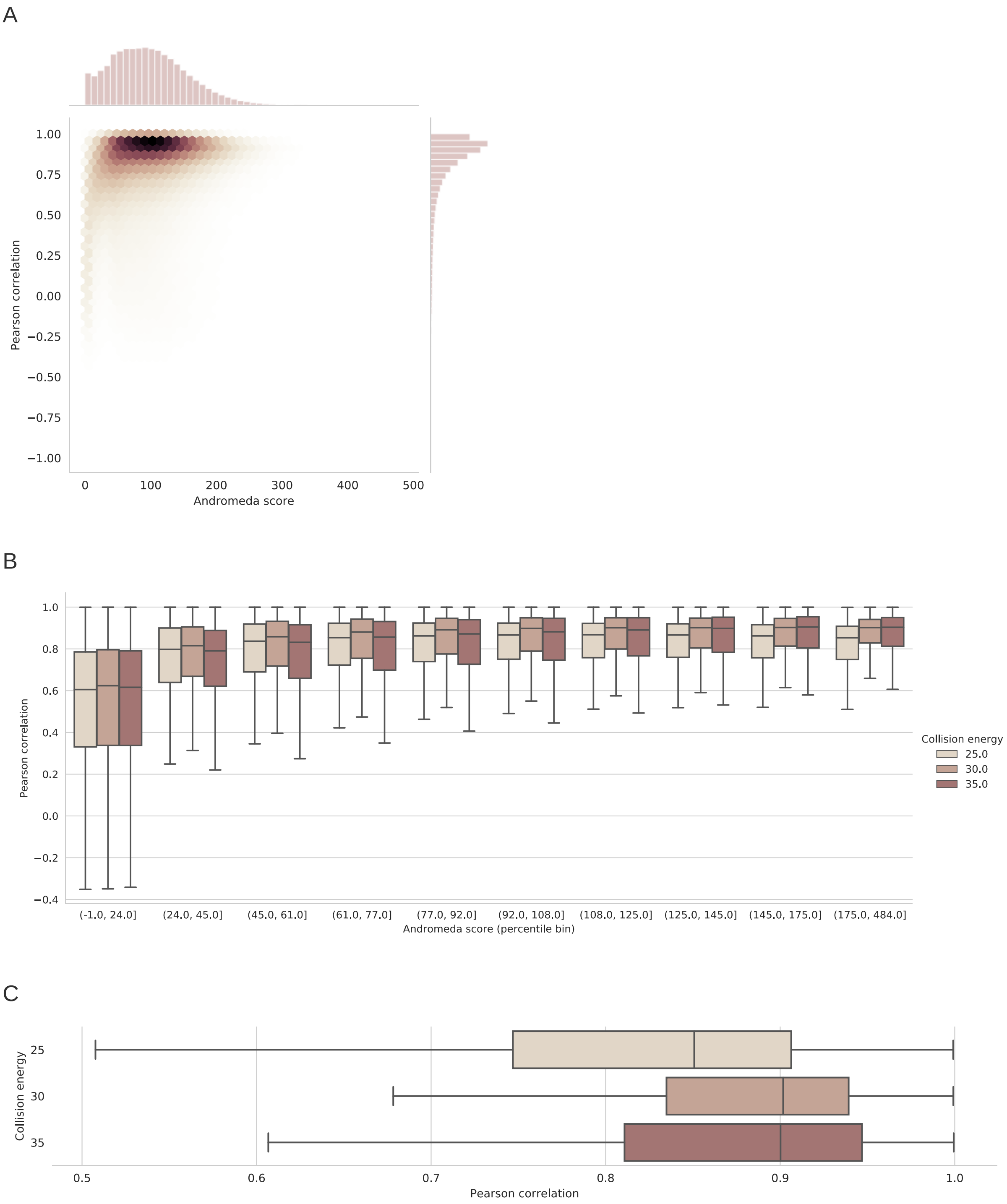


**Supplementary Figure 3.** HCD model evaluation on ProteomeTools synthetic peptide spectra (Zolg et al., 2017, 10.1038/nmeth.4153) across different collision energies (CE). Raw files and MaxQuant identifications were downloaded from PRIDE Archive (PXD004732) for all "3xHCD" MS runs. As no target-decoy strategy was included in the submitted MaxQuant results, we predicted MS²PIP spectra and calculated Pearson correlation coefficients for all MaxQuant identifications and took the Andromeda scores into account in these plots.

A) Two-dimensional histogram, or “Hexbin plot”, (center) and histograms (top and right) of the Andromeda score and Pearson correlation coefficients between MS²PIP predicted and experimental spectra for all included CEs.

B) Boxplots of the Pearson correlation coefficients between MS²PIP predicted and experimental spectra across ten Andromeda score percentiles and split by CE. Every percentile bin contains 10% of the data.

C) Boxplots of the Pearson correlation coefficients between MS²PIP predicted and experimental spectra for all PSMs with an Andromeda score higher than 200, split by CE.
